# The combination of lipopolysaccharide and D-galactosamine administration show positive genotoxic effect in mice liver

**DOI:** 10.1101/2020.01.22.915108

**Authors:** Wenjing Dong, Erqun Song, Yang Song

## Abstract

Lipopolysaccharide (LPS)/D-galactosamine (D-GalN) co-administration induced acute liver injury (ALI) and hepatic fibrosis have been extensively studied. However, whether LPS/D-GalN show genotoxic effect is current unknown. Male mice were divided into eight groups and each group contain eight animals. For the acute administration of LPS/D-GalN, the mice were given a single intraperitoneal (*i.p.*) injection of LPS/D-GalN (25 μg/kg + 250 mg/kg, 25 μg/kg + 500 mg/kg, 50 μg/kg + 500 mg/kg body weight) for 6 h, respectively. The chronic administration was conduct by the *i.p.* injection of LPS/D-GalN (10 μg/kg + 100 mg/kg) every other day for 8 weeks. Saline solution (0.9%) and cyclophosphamide (CTX) (50 mg/kg body weight) injection were used as negative and positive control, respectively. Using single cell gel electrophoresis (SCGE) assay, we found that the acute administration of LPS/D-GalN induces severe DNA damage in mice hepatic cells, but not in brain, sperm and bone marrow cells, implied the genotoxicity of LPS/D-GalN. Interestingly, the chronic treatment of LPS/D-GalN causes significant genotoxic effect in both hepatic and brain cells, but not sperm and bone marrow cells. Histopathological examination in liver and brain section consistent with SCGE results, accordingly. Our study, for the first time, reported the genotoxic potential of LPS/D-GalN co-administration. In addition, LPS/D-GalN administration may serve as an experimental model for further genotoxic study.

## Introduction

Lipopolysaccharide (LPS) is the main endotoxin component of the cell wall of gram-negative bacteria, which play an important role in endotoxic injury [1]. D-galactosamine (D-GalN) is an amino sugar selectively metabolized by the hepatocyte that deplete uridine nucleotides in liver cells and inhibit macromolecule synthesis in hepatocytes [2]. Liver is the main target for LPS challenge, D-GalN significant increases the lethal effect of LPS. Therefore, the acute injection of LPS and D-GalN is a widely used experimental model for acute liver injury (ALI) [3], whilst the long term, low-dose injection of LPS and D-GalN provoke chronic inflammatory responses that resemble hepatic fibrosis [4].

It is widely accepted that LPS/D-GalN-induced hepatotoxicity is associated with excess production of pro-inflammatory cytokines, *e.g.*, tumor necrosis factor-α (TNF-α), interleukin-1β (IL-1β) and interleukin-6 (IL-6), along with the generation of reactive oxygen species (ROS) and the activation of downstream signaling cascade [5, 6]. In fact, ROS may act as the initiator of pro-inflammatory cascades [7]. Therefore, it is not surprising that antioxidants and free radical scavengers were used in the treatment of ALI or hepatic fibrosis.

Beside pro-inflammatory responses, these active free radicals are readily reactive with DNA, which strongly implicate the genotoxic potential of LPS. Previously, a few reports indicated that LPS challenge result in genotoxicity in human peripheral blood mononuclear cells, pre-implanting embryonic and uterine cells [8-10]. Since the combination treatment of LPS and D-GalN amplify the amount of radical production, the genotoxic effect of LPS/D-GalN co-administration ought to be discussed.

Cyclophosphamide (CTX), a commonly used anticancer alkylating agent, is converted into its active metabolites, *e.g.*, acrolein and phosphamide mustard by the hepatic microsomal enzyme. These metabolites are responsible for ROS accumulation and DNA damage [11]. Many studies have been adopted to against CTX-induced genotoxicity.

Here, we hypothesized that LPS/D-GalN treatment is associated with ROS-driven genotoxicity. To address our hypothesis, we then determined the genotoxic potential of LPS/D-GalN in mice models. Integrity of genome was evaluated in the alkaline single cell gel electrophoresis (SCGE) assay. Since the acute and chronic treatment of LPS/D-GalN represent different pathologic patterns, we therefore used two different exposure dosages and durations.

## Materials and methods

### Chemical

LPS (CAS number: 326589-90-6), D-GalN (CAS number: 1772-03-8) and CTX (CAS number: 50-18-0) were purchased from Aladdin Reagent Database Inc. (Shanghai, China). Ethidium bromide (EB) and agarose (normal and low melting point) were purchased from Dingguo Biotechnology Co. (Beijing, China). All other reagents reached analysis level.

### Animal treatment

Male Kunming mice weighing 18-22 g were purchased from Chongqing Academy of Chinese Materia Medica and the following experiments were performed in accordance with the guideline of the Animal Care Committee of Southwest University. Animals were housed in plastic cages under standard conditions (humidity, 50%; temperature, 22 ± 2°C) in a 12 h dark and light cycle, with free access to food and water, in accordance with the guideline of the Animal Care Committee of Southwest University. Mice were acclimatized for at least 7 days before experiments. Mice were randomly divided into 8 groups and each group contain eight animals. For the acute administration of LPS/D-GalN, the mice were given a single intraperitoneal (i.p.) injection of LPS/D-GalN (25 μg/kg + 250 mg/kg, 25 μg/kg + 500 mg/kg, 50 μg/kg + 500 mg/kg body weight) for 6 h, respectively. The chronic administration was conduct by the i.p. injection of LPS/D-GalN (10 μg/kg + 100 mg/kg) every other day for 8 weeks. Saline solution (0.9%) and cyclophosphamide (CTX) (50 mg/kg body weight) injection were used as negative and positive control, respectively. After the end of treatment, mice were anesthetized with isoflurane and sacrificed with regard for alleviation of suffering.

### SCGE assay

After scarifies, the liver, brain, sperm and bone marrow cells in mice were extracted for the following experiments. Cells were diluted to about 1×10^6^ cells/mL with PBS and refrigerated at 4°C. 100 µL normal-melting agarose was placed on single surface grinding board at 4°C for 40 min to cool and curdle. The 50 µL cell suspension and 50 µL low-melting agarose were mixed and spread on glass slides for 40 min at 4°C. Finally, the sides were covered with 100 µL low-melting agarose for 20 min at 4°C. The slides were immersed in lysing solution (2.5 M NaCl, 100 mM Na_2_EDTA, 10 mM Tris, pH 10, and 10% DMSO with 1% Triton X-100) at darkness in 4°C for 1 h. The slides were immersed in a cold alkaline solution (1 mM Na_2_EDTA and 300 mM NaOH, pH>13) for 30 min to spin DNA. Thereafter, electrophoresis was performed for 30 min. The cells were neutralized by neutralizing fluid (0.4 M Tris, pH 7.5), and stained by 10 µg/mL ethidium bromide. The comet’s trailing image was captured by a fluorescence microscope (OLYMPUS IX71). Fifty cells from each of three independent experiments were analyzed with Comet Assay Software Project (CASP) 1.2.2. The tail length (measured from the right edge of the comet head), tail DNA percentage and tail moment (Tail length multiple tail DNA percentage) represent the degree of DNA damage.

### Histopathological examination

Histopathological examination was used to test tissue damage. The whole mice brain and liver of the pretreated mice were obtained, and tissues were soaked in 4% paraformaldehyde and buried by paraffin bag. The fixed brain and liver were cut into 5 μm sections. After hematoxylin-eosin (HE) staining, slides were observed with a fluorescence microscope (TE2000, Nikon, Japan).

### Statistical analysis

The results were expressed as mean ± standard deviations (SD). The data was taken from three independent experiments, and the statistical significance of differences was evaluated by a one-way ANOVA and followed by least significance difference (LSD) multiple comparison tests. P<0.05 was represented statistically significant.

## Results

Histopathological examination was used to detect the damage of mice liver. When compared with the control group, acute injection of LPS/D-GalN in the tested concentrations induced liver cells necrosis, and the necrotic cells expanded from the blood vessels to the entire liver with the increase of LPS/D-GalN concentrations, similar phenomena were shown in the group of CTX (**Figure 1A**). The chronic treatment of LPS/D-GalN caused fibrogenesis feature in the liver of mice (**Figure 1B**). These results suggested the successfully establishment of ALI and hepatic fibrosis models, respectively.

**Figure 1.**
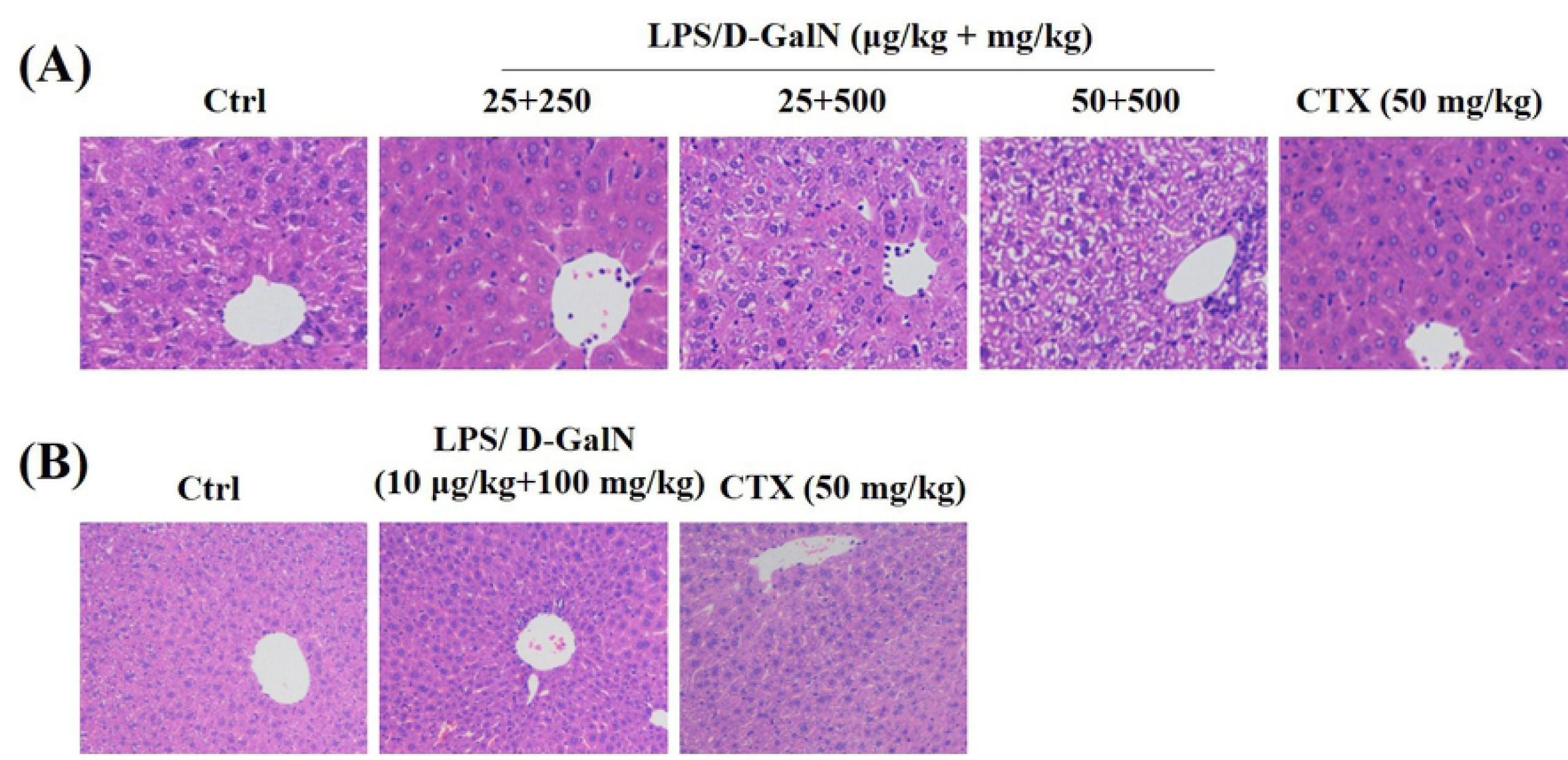
HE staining was used to detect the liver damage. (A) Acute treatment. Mice were injected with LPS/D-GalN (25 μg/kg + 250 mg/kg, 25 μg/kg + 500 mg/kg and 50 μg/kg + 500 mg/kg body weight) for 6 h. HE staining was used to show the lesion of the liver in mice. (B) Chronic treatment. Mice were injected with LPS/D-GalN (10 μg/kg + 100 mg/kg body weight, inject every other day for 8 weeks). HE staining was used to show the lesion of the liver in mice. Saline solution (0.9%) and cyclophosphamide (CTX) (50 mg/kg body weight) injection were used as negative and positive control, respectively.

Oxidative DNA damage includes base and sugar lesions, single or double-strand breaks, DNA-protein or DNA-DNA crosslinks, abasic sites and other exocyclic DNA adducts [12]. Herein, we use single cell gel electrophoresis (SCGE) assay to detect DNA damage occurred in LPS/D-GalN challenged mice. We found that the acute injection of LPS/D-GalN result in concentration-dependent increases of tail moment, tail DNA (%) and olive tail moment in mice hepatic cells (**Table 1** and **Figure 2**).

**Table 1.** Mice were injected with LPS/D-GalN (25 μg/kg + 250 mg/kg, 25 μg/kg + 500 mg/kg and 50 μg/kg + 500 mg/kg body weight) for 6 h. Saline solution (0.9%) and cyclophosphamide (CTX) (50 mg/kg body weight) injection were used as negative and positive control, respectively. SCGE assay was performed. Data were presented as means ± SD, *P < 0.05, **P < 0.01 and ***P < 0.001 indicate significant difference from control group.

|  |  |  | Tail DNA (%) | Tail moment | Olive tail moment |
| --- | --- | --- | --- | --- | --- |
| Liver cell | Ctrl | 0.9% NaCl | 3.61±0.51 | 1.41±0.24 | 2.17±0.25 |
|  | LPS/D-GalN | 25 µg/kg+ 250 mg/kg | 7.19±0.42** | 3.66±0.27 | 4.04±0.37 |
|  |  | 25 µg/kg+500 mg/kg | 7.80±0.43** | 4.46±0.75 | 4.69±0.59* |
|  |  | 50 µg/kg+500 mg/kg | 9.94±0.53*** | 7.50±0.87*** | 6.22±0.46*** |
|  | CTX | 50 mg/kg | 13.12±1.37*** | 11.0±0.27*** | 8.18±1.12*** |
| Brain cell | Ctrl | 0.9% NaCl | 7.66±0.87 | 3.31±0.51 | 1.34±0.21 |
|  | LPS/D-GalN | 25 µg/kg+ 250 mg/kg | 9.11±0.84 | 3.78±0.50 | 0.74±0.12 |
|  |  | 25 µg/kg+500 mg/kg | 8.89±0.97 | 4.31±0.68 | 0.59±0.10 |
|  |  | 50 µg/kg+500 mg/kg | 6.64±1.10 | 4.71±1.02 | 1.79±0.32 |
|  | CTX | 50 mg/kg | 23.20±1.62*** | 22.59±2.01*** | 7.69±0.80*** |
| Bone marrow cell | Ctrl | 0.9% NaCl | 1.39±0.16 | 0.32±0.06 | 0.72±0.11 |
|  | LPS/D-GalN | 25 µg/kg+ 250 mg/kg | 2.68±0.29 | 0.83±0.14 | 1.30±0.15 |
|  |  | 25 µg/kg+500 mg/kg | 1.80±0.22 | 0.40±0.09 | 0.85±0.11 |
|  |  | 50 µg/kg+500 mg/kg | 2.22±0.26 | 1.14±0.21 | 1.45±0.12 |
|  | CTX | 50 mg/kg | 9.70±1.14*** | 6.72±1.21*** | 6.77±0.91*** |
| Sperm cell | Ctrl | 0.9% NaCl | 2.76±0.62 | 0.81±0.28 | 0.66±0.14 |
|  | LPS/D-GalN | 25 µg/kg+ 250 mg/kg | 0.89±0.12 | 0.24±0.16 | 0.27±0.09 |
|  |  | 25 µg/kg+500 mg/kg | 2.10±0.30 | 0.57±0.12 | 0.62±0.09 |
|  |  | 50 µg/kg+500 mg/kg | 2.90±0.26 | 0.47±0.05 | 0.84±0.10 |
|  | CTX | 50 mg/kg | 17.38±8.11*** | 8.20±0.96*** | 5.1±0.28*** |

**Figure 2.**
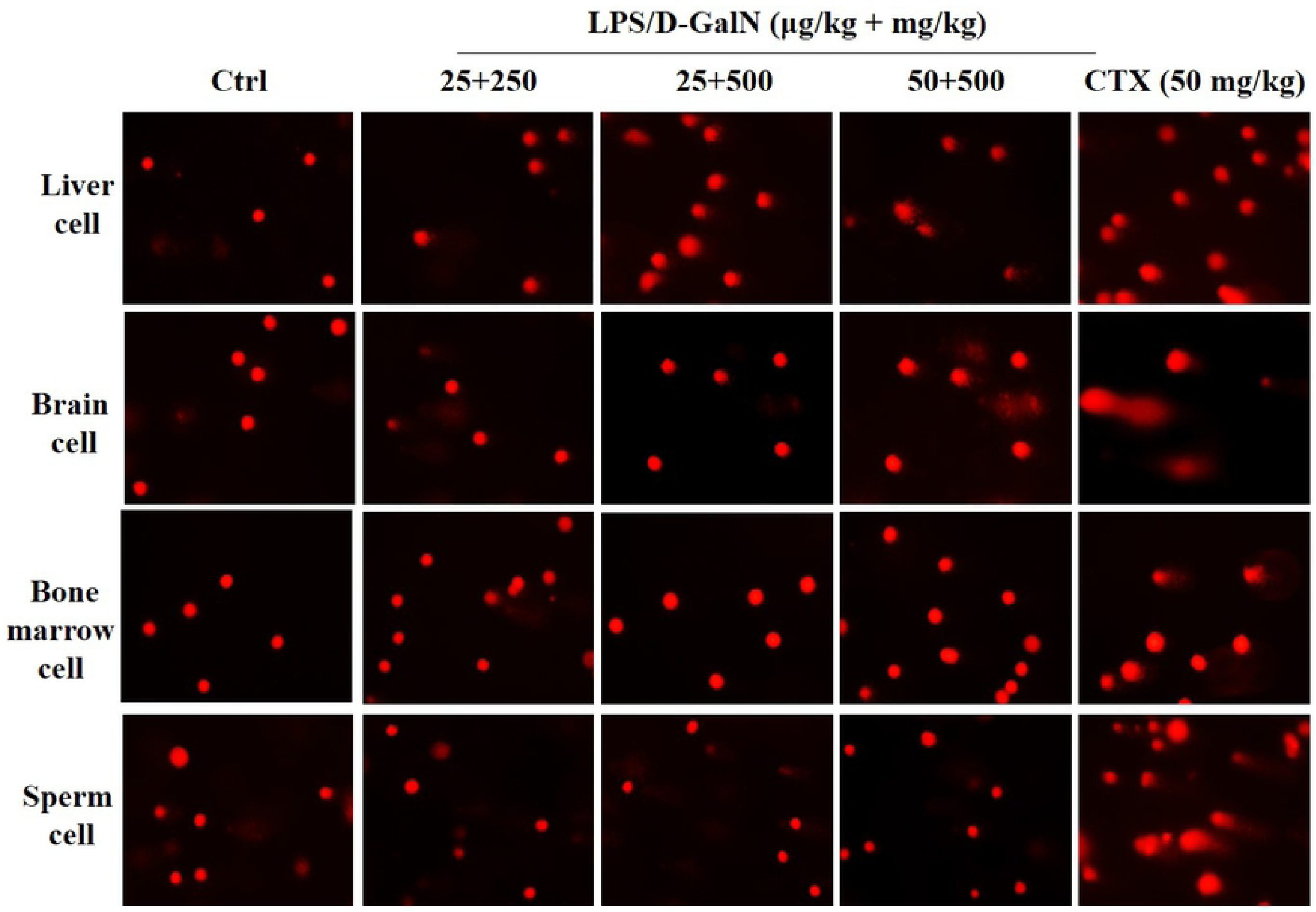
Mice were injected with LPS/D-GalN (25 μg/kg + 250 mg/kg, 25 μg/kg + 500 mg/kg and 50 μg/kg + 500 mg/kg body weight) for 6 h. Saline solution (0.9%) and cyclophosphamide (CTX) (50 mg/kg body weight) injection were used as negative and positive control, respectively. SCGE assay was performed. The comet’s trailing image was captured by a fluorescence microscope (OLYMPUS IX71).

Previous studies illustrated that administration of LPS in mice induced inflammation in brain [13], testicles [14] and bone marrow [15]. Next, we investigate whether LPS/D-GalN showed similar effect in these targets. However, there were no significant DNA damage in brain, sperm and bone marrow cells upon the acute injection of LPS/D-GalN (**Table 1**). CTX was used as positive control to verify the credibility of our experimental results. Surprisingly, CTX showed positive genotoxic effect not only in hepatic cells, but also other three cells, which consistent with previous studies [16-19]. This data implied the different toxicokinetics between CTX and LPS/D-GalN.

Although liver is the major target for LPS/D-GalN challenge, the pro-inflammatory cytokines may release from the primary organ and resulting in chronic systemic inflammation. It is current unknown whether chronic LPS/D-GalN challenge induces ROS propagation through liver to other organs. Surprisingly, compare to the negative control, the chronic combined application of LPS and D-GalN induced clear DNA migration in mice liver and brain cells, statistical significances of tail moment, tail DNA (%) and olive tail moment were found in both cells, respectively (**Table 2** and **Figure 3**).

**Table 2.**
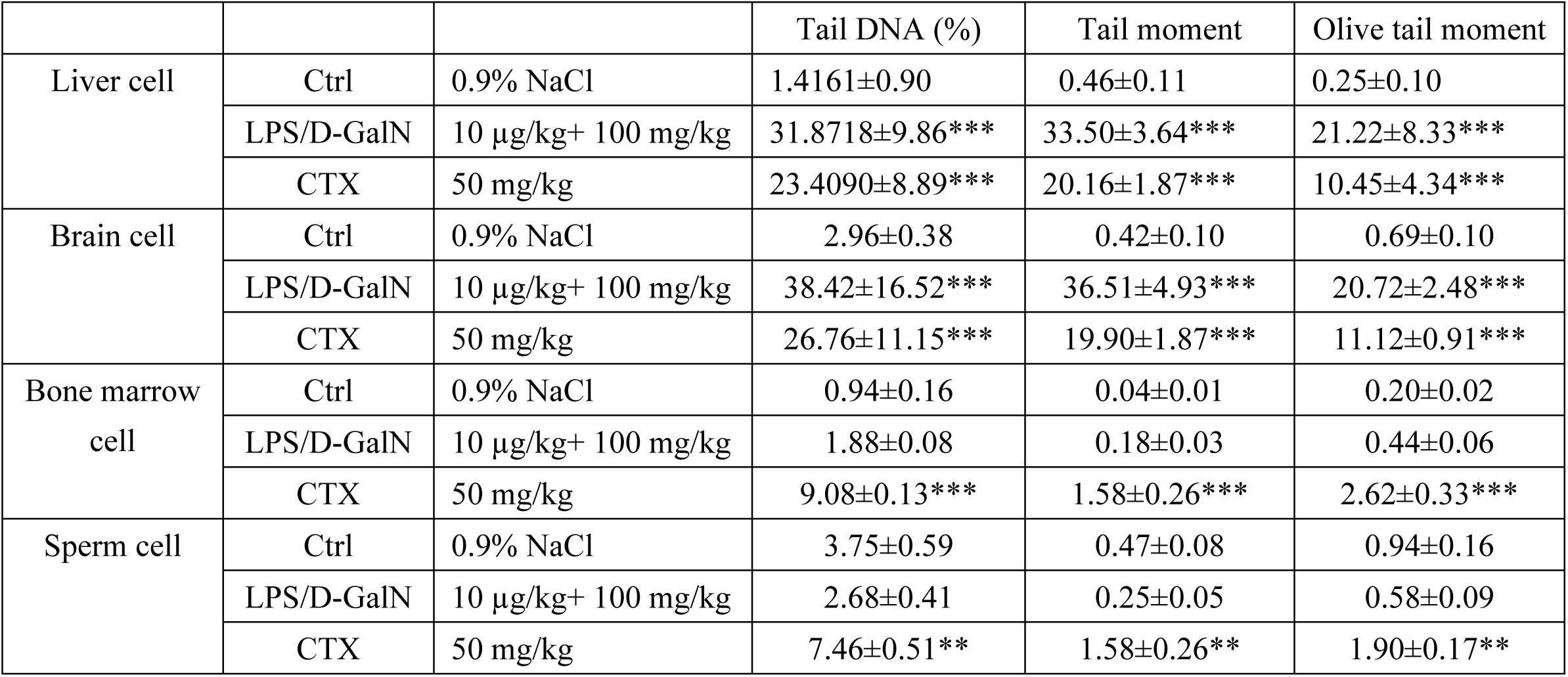
Mice were injected with LPS/D-GalN (10 μg/kg + 100 mg/kg body weight, inject every other day for 8 weeks). Saline solution (0.9%) and cyclophosphamide (CTX) (50 mg/kg body weight) injection were used as negative and positive control, respectively. SCGE assay was performed. Data were presented as means ± SD, *P < 0.05, **P < 0.01 and ***P < 0.001 indicate significant difference from control group.

|  |  |  | Tail DNA (%) | Tail moment | Olive tail moment |
| --- | --- | --- | --- | --- | --- |
| Liver cell | Ctrl | 0.9% NaCl | 1.4161±0.90 | 0.46±0.11 | 0.25±0.10 |
|  | LPS/D-GalN | 10 µg/kg+ 100 mg/kg | 31.8718±9.86*** | 33.50±3.64*** | 21.22±8.33*** |
|  | CTX | 50 mg/kg | 23.4090±8.89*** | 20.16±1.87*** | 10.45±4.34*** |
| Brain cell | Ctrl | 0.9% NaCl | 2.96±0.38 | 0.42±0.10 | 0.69±0.10 |
|  | LPS/D-GalN | 10 µg/kg+ 100 mg/kg | 38.42±16.52*** | 36.51±4.93*** | 20.72±2.48*** |
|  | CTX | 50 mg/kg | 26.76±11.15*** | 19.90±1.87*** | 11.12±0.91*** |
| Bone marrow cell | Ctrl | 0.9% NaCl | 0.94±0.16 | 0.04±0.01 | 0.20±0.02 |
|  | LPS/D-GalN | 10 µg/kg+ 100 mg/kg | 1.88±0.08 | 0.18±0.03 | 0.44±0.06 |
|  | CTX | 50 mg/kg | 9.08±0.13*** | 1.58±0.26*** | 2.62±0.33*** |
| Sperm cell | Ctrl | 0.9% NaCl | 3.75±0.59 | 0.47±0.08 | 0.94±0.16 |
|  | LPS/D-GalN | 10 µg/kg+ 100 mg/kg | 2.68±0.41 | 0.25±0.05 | 0.58±0.09 |
|  | CTX | 50 mg/kg | 7.46±0.51** | 1.58±0.26** | 1.90±0.17** |

**Figure 3.**
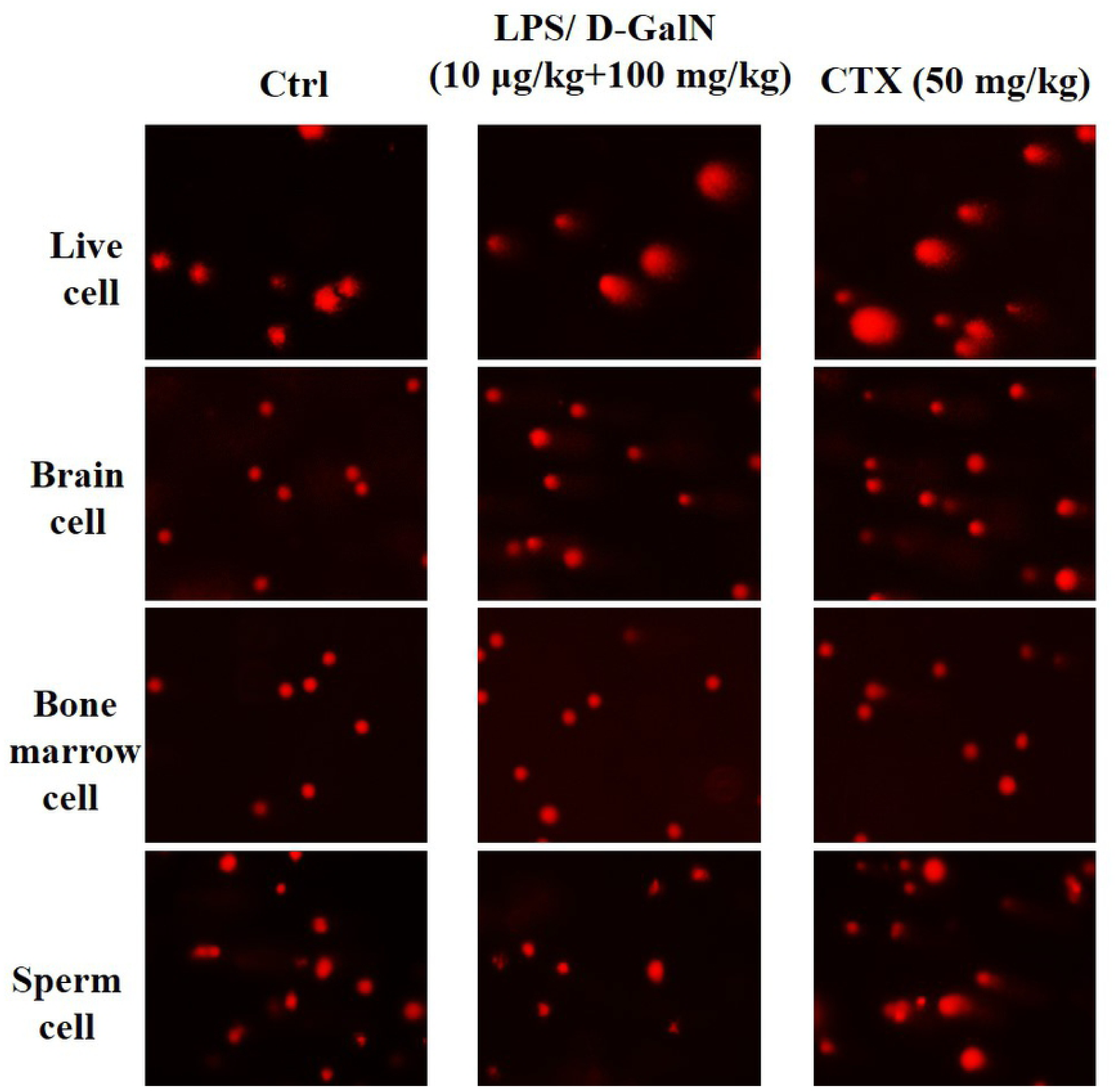
Mice were injected with LPS/D-GalN (10 μg/kg + 100 mg/kg body weight, inject every other day for 8 weeks). Saline solution (0.9%) and cyclophosphamide (CTX) (50 mg/kg body weight) injection were used as negative and positive control, respectively. SCGE assay was performed. The comet’s trailing image was captured by a fluorescence microscope (OLYMPUS IX71).

Histopathological examination in brain section are consistent with SCGE result. Chronic administration of LPS/D-GalN induces wrinkled cerebral cortex neurons, whilst there was no significant change in the cerebral cortex of mice with acute LPS/D-GalN administration (**Figure 4**).

**Figure 4.**
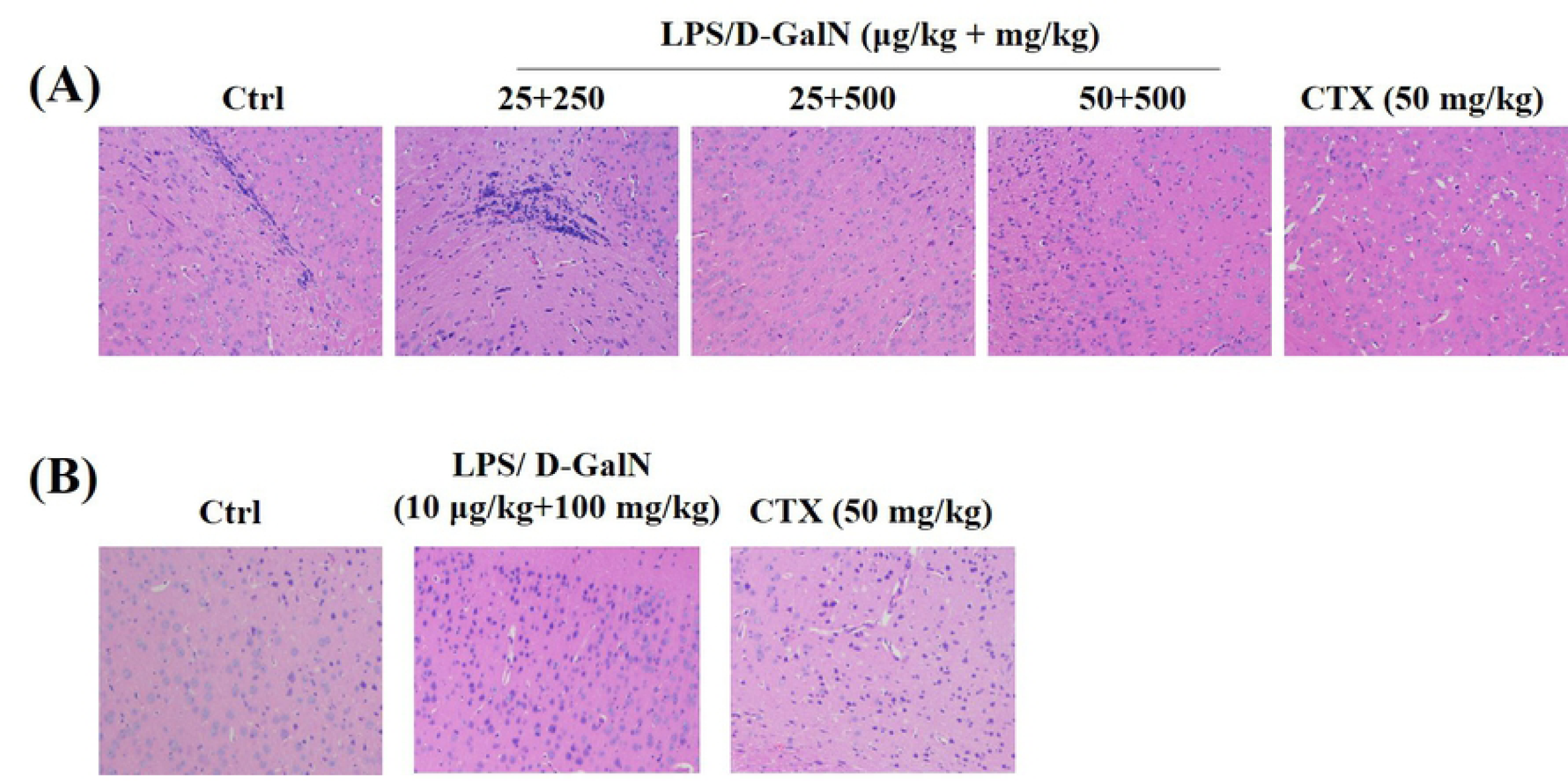
HE staining was used to detect the brain damage. **(A)** Mice were injected with LPS/D-GalN (25 μg/kg + 250 mg/kg, 25 μg/kg + 500 mg/kg and 50 μg/kg + 500 mg/kg body weight) for 6 h. Saline solution (0.9%) and cyclophosphamide (CTX) (50 mg/kg body weight) injection were used as negative and positive control, respectively. **(B)** Mice were injected with LPS/D-GalN (10 μg/kg + 100 mg/kg body weight, inject every other day for 8 weeks). Saline solution (0.9%) and cyclophosphamide (CTX) (50 mg/kg body weight) injection were used as negative and positive control, respectively.

On the contrary, there were not obviously increased DNA damage in bone marrow and sperm cells after chronic LPS/D-GalN administration (**Table 2**). CTX showed strong genotoxic effect in four cell lines. These result suggested that LPS/D-GalN activated a ROS-mediated toxic accumulation process in the brain after a long-term administration; however, exact molecular mechanism needs further investigation.

## Discussion

LPS is known to mimic the pathophysiology of gram-negative bacterial infections, the combined treatment of D-GalN is a well-established model of liver injury in human and other mammals. The current approach for the first time reported that the co-administration of LPS/D-GalN show strong genotoxic effect in their primary target, liver. The co-administration of LPS/D-GalN resulted in a significant increase in Tail DNA (%), tail moment and olive tail moment tail moment in exposed animal livers, indicating that LPS/D-GalN is genotoxic. Our findings are important because they suggested that the combination of LPS/D-GalN administration may serve as an experimental model for genotoxic study.

Although liver is the major target for LPS insult, LPS-associated damages have been investigated in different organs, for instance, lung, heart and kidney [20-22]. Our present study conclude that the acute injection of LPS/D-GalN induces positive genotoxic effect only in liver cells, whilst the chronic injection of LPS/D-GalN induces positive genotoxic effect both in liver and brain cells. Neither acute nor chronic injection of LPS/D-GalN induce significant genotoxic effect in bone marrow or sperm cells. Please note the bio-distribution of LPS/D-GalN has not been discussed in this study, the accumulation of LPS/D-GalN is probably not sufficient to elicit a genotoxic effect in the brain cells of acute injection model. In addition, our data also reflect the dissimilar sensitivity of between tissues in responses to LPS/D-GalN challenge.

Brain inflammation is associated with many age-related central nervous system diseases, for example, Parkinson’s and Alzheimer’s diseases. Although there is little likelihood that brain suffer direct endotoxin insult, pro-inflammatory neurodegeneration in brain can be induced by head trauma, ischemia, and protein aggregates of amyloid β, which are commonly leading to perpetuated release of neurotoxic cytokines. The chronic but not acute LPS/D-GalN treatment resulted in the positive genotoxicity in brain cells, implied that brain may not the primary target for LPS/D-GalN. Alternatively, the release of pro-inflammatory cytokines in the liver spread through blood circulation and thus leading to systemic inflammation. Brain is a relative vulnerable organ and perpetuated release of neurotoxic cytokines and causes significant genotoxic effect in brain cells.

LPS is not only endogenous but also exogenous toxin that generates free radical, *e.g.*, hydroxyl and nitric oxide radical [23]. Thereafter, the principal toxic mechanism of LPS is known to cause damage to protein, DNA and lipids. Continuously high ROS level result in permanent DNA damage, genome instability, and in turn cancer initiation and progression. However, other signaling pathways also participated in LPS-induced toxicities, for instance, inflammation, autophagy, and *etc*. Inflammatory responses also play major roles in LPS-induced toxic effect. Specifically, Toll like receptor 4 (TLR4) is the main receptor of LPS, that leads to inflammation through TLR4/nuclear factor kappa B (NF-κB) signaling. Next, the activation of NF-κB cascade leads to the release of inflammatory cytokines. D-GalN co-administration further amplifies ROS production. In this regard, LPS/D-GalN administration was previously believed in the evokes of complication of liver sepsis. Here, using SCGE assay, we first demonstrated that LPS/D-GalN challenge showed the occurrence of DNA damage *in vivo*. In support of our current result, Verma *et al*. have also found out tissue inflammation and genotoxicity in mice following LPS challenge [24]. Accumulating evidences indicated that inflammation is a protective response against ROS attack. In addition, inflammation is also a promotor of repair and regeneration in injured cells [25]. Interestingly, autophagic response activation by LPS stimulation are pervasive. Autophagy is a homeostatic process, which is responsible for ameliorate LPS-induced damages or inflammatory status. It is proposed that autophagic degradation of damaged proteins or other biomarcomolecules that play an essential role on cellular homeostasis. Xanthine oxidase and heme oxygenase were also found to be activated, implied the chromosome damaging properties of LPS [26]. Together, inherent adaptive mechanisms, including inflammation and autophagy, serves protective effect on LPS-induced damages, however, whether and how inflammation and autophagy-related molecules, along with pro-death signaling (*e.g.*, apoptosis, necroptosis and necrosis), communicate with genotoxic insults need further investigation. Thus, further studies are also needed for discuss this possibility.

In a general perspective, enzymatic antioxidants (*e.g.*, superoxide dismutase, catalase, glutathione peroxidase and glutathione reductase) or non-enzymatic antioxidants (*e.g.*, glutathione, N-Acetyl-L-cysteine and selenium) play essential role in protecting cells from ROS insult through the elimination of ROS propagation. Indeed, numerous of studies were conducted to screening anti-inflammatory drugs or compounds in LPS model. It is well known that the biologic consequences of LPS challenge include the release of cytokines. However, to the best of our knowledge, there has no information regarding to the use of LPS/D-GalN in genotoxic model. Besides the subsequent elucidation of anti-inflammatory parameters, this study can provide information on potential therapeutics for both chronic and acute genotoxic processes. Of note, the fundamental principle of LPS/D-GalN induced liver (and brain) damage is belong to ROS explosion, no matter which endpoint were investigated.

In conclusion, we conclude that acute and chronic exposure to LPS/D-GalN results in genotoxicity and suggest the using of LPS/D-GalN co-administration as an experimental model/positive control of hepatic genotoxicity for further studies.

## Data Availability

All relevant data are within the paper.

